# Triple A patient cells suffering from mitotic defects fail to localize PGRMC1 to mitotic kinetochore fibers

**DOI:** 10.1101/381541

**Authors:** Ramona Jühlen, Dana Landgraf, Angela Huebner, Katrin Koehler

## Abstract

- Investigating cell division in human adrenal cells we show that proliferation is decreased upon overexpression of ALADIN, PGRMC1 or PGRMC2.
- In immunofluorescence experiments using human adrenal cells and triple A patient fibroblasts we observed that during cell division PGRMC1 localizes to the microtubule kinetochore-fibers in metaphase and to the mid-body in telophase.
- Depletion of ALADIN results in mis-localization of Aurora A and PGRMC1 in metaphase cells of the human adrenal cell line and fibroblasts derived from patients with triple A syndrome.
- In real time PCR using RNA of fibroblasts of triple A syndrome patients and healthy controls we measured an increased expression of PGRMC2 in cells with ALADIN mis-function compared to the control cells.
- We hypothesize that a loss of the regulatory interaction between ALADIN and PGRMC2 leads to an over-regulation and over-expression of PGRMC2 and displaces PGRMC1 at the metaphase spindle. This diminishing of PGRMC1 concentration at kinetochore fibers may lead to mitotic errors and pro - liferation arrest.

**ABSTRACT:** Membrane-associated progesterone receptors are restricted to the endoplasmic reticulum and are shown to regulate the activity of cytochrome P450 enzymes which are involved in steroidogenesis or drug detoxification. PGRMC1 and PGRMC2 belong to this group of microsomal receptors and are of interest due to their suspected role during cell cycle. PGRMC1 and PGRMC2 are thought to bind to each other thereby suppressing entry into mitosis. We could previously report that PGRMC2 interacts with the nucleoporin ALADIN which when mutated results in the autosomal recessive disorder triple A syndrome. ALADIN is a novel regulator of mitotic controller Aurora kinase A and depletion of this nucleoporin leads to microtubule instability. In the current study, we present that proliferation is decreased when ALADIN, PGRMC1 or PGRMC2 are over-expressed. Furthermore, we find that depletion of ALADIN results in mis-localization of Aurora kinase A and PGRMC1 in metaphase cells. Additionally, PGRMC2 is over-expressed in triple A patient fibroblasts. Our results emphasize the possibility that loss of the regulatory interaction between ALADIN and PGRMC2 gives rise to a depletion of PGRMC1 at kinetochore fibers and to mitotic errors. This observation may explain part of the symptoms seen in triple A syndrome patients.

## INTRODUCTION

ALADIN is a scaffold nucleoporin (NUP) anchored within the nuclear pore complex by the transmembrane NUP NDC1 (Kind *et al*., 2009; Yamazumi *et al*., 2009). ALADIN seems to be involved in building the structural scaffold backbone of the nuclear pore complex (Rabut *et al*., 2004). Over the last years it has been shown that nuclear pore complexes and its NUPs have fundamental functions beyond nucleo-cytoplasmic transport and control cellular dysfunction in a variety of cellular pathways, especially during mitosis (Fahrenkrog, 2014; Nofrini *et al*., 2016; Sakuma and D’Angelo, 2017). The first report that ALADIN plays a role during cell division was published in 2015 (Carvalhal *et al*., 2015). In cooperation with Carvalhal et al. we proposed ALADIN as novel regulator of mitotic kinase Aurora kinase A and showed that depletion of the nucleoporin resulted in impaired mitotic spindle assembly and chromosomal alignment at the metaphase plate (Carvalhal *et al*., 2015). Furthermore, we could recently document that ALADIN is necessary for murine oocyte maturation and for specific stages during meiosis (Carvalhal *et al*., 2017).

Mutations in the human *AAAS* gene, coding for the protein ALADIN, lead to the autosomal recessive disorder named triple A syndrome (Tullio-Pelet *et al*., 2000; Handschug *et al*., 2001). Triple A patients present with three distinct symptoms: absent adrenal glucocorticoid and mineralocorticoid synthesis (adrenal insufficiency), impaired movement of the stomach cardia (achalasia) and loss of tear production (alacrima) (Allgrove *et al*., 1978). These symptoms are highly heterogeneous and are accompanied by progressive impairments of the central, peripheral or autonomous nervous system (Allgrove *et al*., 1978). Most mutations in *AAAS* result in a mis-localization of ALADIN to the cytoplasm (Cronshaw and Matunis, 2003; Krumbholz *et al*., 2006).

Previously, we identified microsomal PGRMC2 as novel interactor for the nucleoporin ALADIN and provided new insights into the molecular function of the nucleoporin in the pathogenesis of triple A syndrome (Jühlen *et al*., 2016). PGRMC2 belongs to the group of membrane-associated progesterone receptors. These receptors are restricted to the endoplasmic reticulum (ER) and are thought to regulate the activity of microsomal cytochrome (CYP) P450 enzymes which are involved in steroidogenesis or drug detoxification (Cahill and Medlock, 2017). The first identified membrane-associated progesterone receptor, PGRMC1, gained wide-spread attention due to its several implications in cancerogenesis (Falkenstein *et al*., 1996; Clark *et al*., 2016; Kabe *et al*., 2016; Ryu *et al*., 2017). The mixed-function oxidase system of CYP P450 enzymes requires a donor transferring electrons from NADPH to reduce the prosthetic heme group (Pandey and Flück, 2013). PGRMC1 and PGRMC2 contain a CYP b5-similar heme-binding domain (Ryu *et al*., 2017). PGRMC1 forms stable protein-protein complexes with CYP51A1, CYP7A1, CYP21A2 and CYP3A4 (Hughes *et al*., 2007). Additionally, PGRMC1 is able to activate CYP19 aromatase (Ahmed *et al*., 2012). PGRMC1 is shown to physiologically affect cholesterol/steroid biosynthesis and metabolism (Ryu *et al*., 2017). It is known that PGRMC2 has similar interaction potential, alters activity of CYP3A4 as possible electron donor, and binds CYP21A2 (Albrecht *et al*., 2012; Wendler and Wehling, 2013). Most recently, both PGRMC1 and PGRMC2 were identified as putative interacting partners of ferrochelatase, an enzyme catalyzing the terminal step in the heme biosynthetic pathway, thereby possibly controlling heme release as chaperone or sensor (Piel *et al*., 2016). Interaction of ALADIN with PGRMC2 at the perinuclear ER could influence CYP P450 enzyme activity through electron transfer from NADPH and/or control heme synthesis. In triple A syndrome, altered CYP P450 enzyme activity would consecutively contribute to adrenal atrophy (Jühlen *et al*., 2016).

Human PGRMC1 and PGRMC2 share 67 % of their protein sequence (Cahill, 2017; Cahill and Medlock, 2017). Deficiency of either PGRMC1 or PGRMC2 decreases the anti-apoptotic and anti-mitotic action of progesterone (Peluso *et al*., 2014). Additionally, increased expression of PGRMC1 or PGRMC2 inhibits entry into cell cycle (Griffin *et al*., 2014; Peluso *et al*., 2014). On the one hand, PGRMC1 is distributed with β- and γ-tubulin to the mitotic bipolar spindle and spindle poles in metaphase cells and on the other hand, with Aurora kinase B in meiotic cells (Luciano *et al*., 2010; Lodde and Peluso, 2011). Furthermore, PGRMC1 is thought to regulate microtubule stability (Griffin *et al*., 2014). PGRMC2 is shown to localize to the mitotic spindles in metaphase and anaphase cells and shall interact with cyclin-dependent kinase 11B (p58) (Griffin *et al*., 2014). PGRMC1 and PGRMC2 are reported to interact and furthermore, to bind to each other during metaphase, thereby suppressing entry into cell cycle (Peluso *et al*., 2014).

Interestingly, in a large scale interactome mapping of the centrosome-cilium interface the nucleoporin ALADIN and both microsomal PGRMC1 and PGRMC2 have been identified to localize to the cilium transition zone (Hanson *et al*., 2014; Gupta *et al*., 2015). The centrosome is an important regulator of cell cycle progression and mitotic spindle assembly (Scholey *et al*., 2003). Furthermore, ALADIN is strongly dephosphorylated during mitotic exit, whereas PGRMC1 and PGRMC2 are phosphorylated during early mitotic exit (Malik *et al*., 2009; McCloy *et al*., 2015). Obviously, ALADIN, PGRMC1 and PGRMC2 seem to have critical roles for the formation and function of the mitotic spindle apparatus in somatic cells.

Here, we report that proliferation in human adrenal cells is decreased after increased expression of ALADIN, PGRMC1 or PGRMC2. We show that PGRMC1 localizes to the microtubule kinetochore-fibers in metaphase and to the midbody in telophase of human adrenal cells and fibroblasts. Further, we present that PGRMC1 and Aurora kinase A are mislocalized in metaphase triple A patient fibroblasts. We observed that patient fibroblasts present with increased expression of PGRMC2 and we hypothesize that a depletion of ALADIN in these cells leads to a dysregulation of PGRMC2 and displaces PGRMC1 at the metaphase spindle.

## RESULTS AND DISCUSSION

### Adrenal cell proliferation is decreased upon over-expression of ALADIN, PGRMC1 or PGRMC2 and down-regulation of ALADIN

In primary skin fibroblasts of triple A patients the cellular proliferation rate is decreased and doubling time of patient cells is significantly increased compared to cells of healthy donors (Kind *et al*., 2010). Additionally, patient cells show features of senescence even though these cells have not reached replicative senescence as it has been documented for normal skin fibroblasts (Lee *et al*., 2002; Kind *et al*., 2010). The nucleoporin ALADIN is ubiquitously expressed with predominant expression in adrenal gland, gastrointestinal and central nervous system structures (Huebner *et al*., 2002). That may be a reason why patients with triple A syndrome present with characteristic pathogenesis in distinct tissues: adrenal insufficiency, achalasia, alacrima and involvement of the central, peripheral and autonomous nervous system (Allgrove *et al*., 1978). In order to address the pathogenesis in adrenal tissue in the patients, we reported that loss of ALADIN leads to an impairment of glucocorticoid and mineralocorticoid synthesis in adrenal cells *in vitro* (Jühlen *et al*., 2015).

To test whether depletion of ALADIN also results in decreased cellular proliferation in adrenal cells we used inducible adrenocortical NCI-H295R1-TR cells stably expressing *AAAS* shRNA (*AAAS* knock-down) and monitored cellular proliferation using live cell imaging for at least 65 h. *AAAS* knock-down resulted in decreased proliferation in adrenal cells (growth constant k (slope of linear regression line) =0.044) compared to control cells expressing a scrambled shRNA (k=0.062) (wild-type cells: k=0.2) (Figure 1A). Surprisingly, in live cell imaging stable over-expression of N-terminal-GFP-tagged ALADIN in adrenocortical NCI-H295R cells also impaired cellular proliferation (k=0.082) compared to over-expression of GFP alone in these cells (k=0.305) (wild-type cells: k=0.16) (Figure 1B). It can be assumed that an equilibrated level of ALADIN protein is prerequisite for successful cellular proliferation and that depletion or accumulation of the nucleoporin impairs cellular proliferation. That hypothesis that an equilibrated level of ALADIN is critical for cell division was already postulated by Carvalhal et al. (Carvalhal *et al*., 2015).

**Figure 1.**
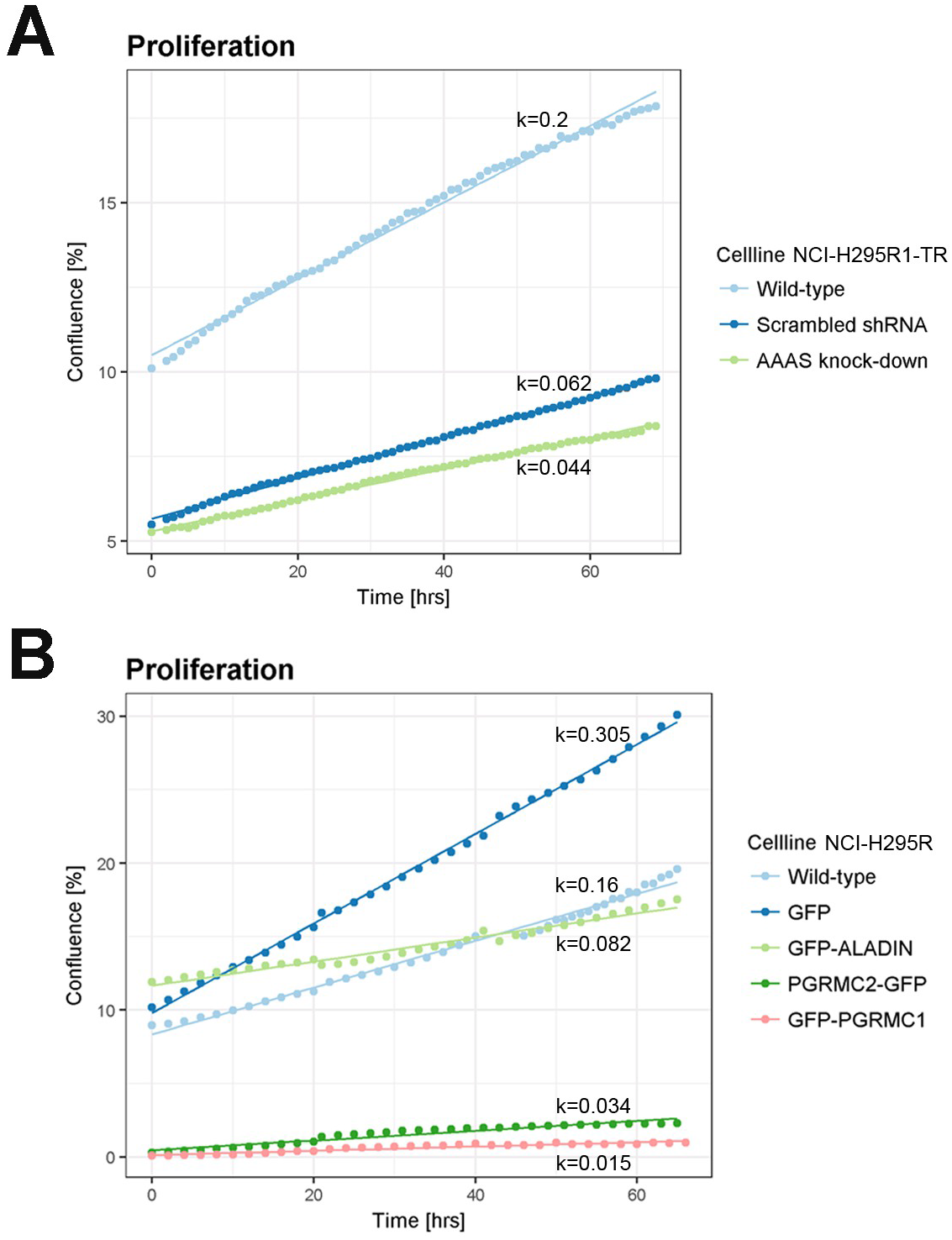
Cellular proliferation is impaired upon over-expression of ALADIN, PGRMC1 or PGRMC2. Equal cell numbers were seeded 48 hours before confluence measurement using live cell imaging on IncuCyte Zoom (Essen BioScience). In case of transient transfection with PGRMC2-GFP and GFP-PGRMC1 the proliferation of fluorescent cells was measured which starts at cell density of about zero. Growth curve analysis and growth constant k (slope of regression curve) calculation was done using multilevel regression technique using R Studio.

We firstly investigated, whether cellular proliferation defects are caused by diminished expression of the nucleoporin ALADIN. Secondly, we wanted to know whether an over-expression of PGRMC2, a novel discovered interactor of the nucleoporin ALADIN, impairs the function during proliferation. In ovarian granulosa cells levels of PGRMC2 decrease during G1 phase of cell cycle (Griffin *et al*., 2014). Increased expression of PGRMC1 or PGRMC2 in these cells is documented to suppress entry into cell cycle (Griffin *et al*., 2014; Peluso *et al*., 2014). During metaphase both proteins are thought to interact and to impair progression of cell cycle (Peluso *et al*., 2014). During live cell imaging we observed that transient overexpression of C-terminal-GFP-tagged PGRMC2 in adrenocortical NCI-H295R cells resulted in decreased cellular proliferation (k=0.034) (Figure 1B). The same phenotype was seen in adrenocortical cells transiently over-expressing N-terminal-GFP-tagged PGRMC1 (k=0.015) (Figure 1B). These results are in accordance with the literature and emphasize a simultaneous role of PGRMC1 and PGRMC2 during cellular proliferation.

### PGRMC1 is restricted to metaphase spindle during mitosis and mid-body during cytokinesis

Next we determined the localization of PGRMC1 and PGRMC2 during cell division in adrenal cells. So far, we saw that over-expression of ALADIN, PGRMC1 or PGRMC2 leads to an impairment of adrenal cellular proliferation (Figure 1B). Thus, we assumed that each of these proteins plays an important role during regulation of cell division. It is known that ALADIN is a novel regulator of mitotic controller Aurora kinase A (Carvalhal *et al*., 2015). Humans have three classes of serine/threonine Aurora kinases: Aurora A, Aurora B and Aurora C (Marumoto *et al*., 2005). The homologous Aurora kinase A and Aurora kinase B are expressed in most cell types. Despite different localization and activation during cell cycle, they both regulate progression through cell cycle from G2 phase to cytokinesis (Marumoto *et al*., 2005). In *Drosophila* ALADIN diffuses loosely through the spindle during mitosis (Carvalhal *et al*., 2015). In HeLa cells ALADIN could be found within the mitotic spindle accumulated at spindle poles but the highest amount appeared between centrosome and metaphase plate (Carvalhal *et al*., 2015). This high concentration of ALADIN protein is thought to be an ER-membrane-associated pool of the nucleoporin since it partially co-localizes with calnexin which is an integral microsomal protein (Carvalhal *et al*., 2015). PGRMC1 and PGRMC2 are documented to localize to the mitotic spindle apparatus in metaphase ovarian granulosa cells but their role at the spindle apparatus and during cell cycle is not known (Lodde and Peluso, 2011; Griffin *et al*., 2014).

Firstly, using immunofluorescence in adrenal NCI-H295R cells we show that PGRMC1 throughout prophase and metaphase localized to the centrosome, bipolar spindle and spindle poles (Figure 2A). During anaphase PGRMC1 diffuses weakly around chromosomes and in telophase before cytokinesis (cell-cell scission) PGRMC1 localizes to the mid-body (Figure 2A). Secondly, during prophase PGRMC1 partially co-localizes with mitotic Aurora kinase A (AURKA) to the centrosome and during metaphase and telophase to spindle poles and mid-body (Figure 2A). During telophase and cell-cell scission PGRMC1 could also be found with mitotic Aurora kinase B (AURKB) at the mid-body (Figure 2B). PGRMC1 has been documented to associate with Aurora kinase B during meiosis but no such role has been assigned during mitosis (Luciano *et al*., 2010). Both Aurora kinase A and Aurora kinase B phosphorylate and thereby regulate a variety of mitotic spindle substrates. Dysregulation of these kinases results in fatal mitotic errors (Marumoto *et al*., 2005). Aurora kinase A has distinct roles in centrosome maturation, entry into mitosis, spindle assembly and microtubule (MT) function during anaphase. Aurora kinase B is essential for bi-orientation of chromosomes and cytokinesis (Barr and Gergely, 2007). It is tempting to assume that PGRMC1 is a substrate being phosphorylated by one or both of these mitotic kinases and indeed, PGRMC1 has been found to be phosphorylated at three postulated serines during early mitotic exit (McCloy *et al*., 2015). Further research should address whether PGRMC1 is phosphorylated by Aurora kinases and should uncover the regulatory effect during mitosis of phosphorylation on one of the three postulated serines in PGRMC1.

**Figure 2.**
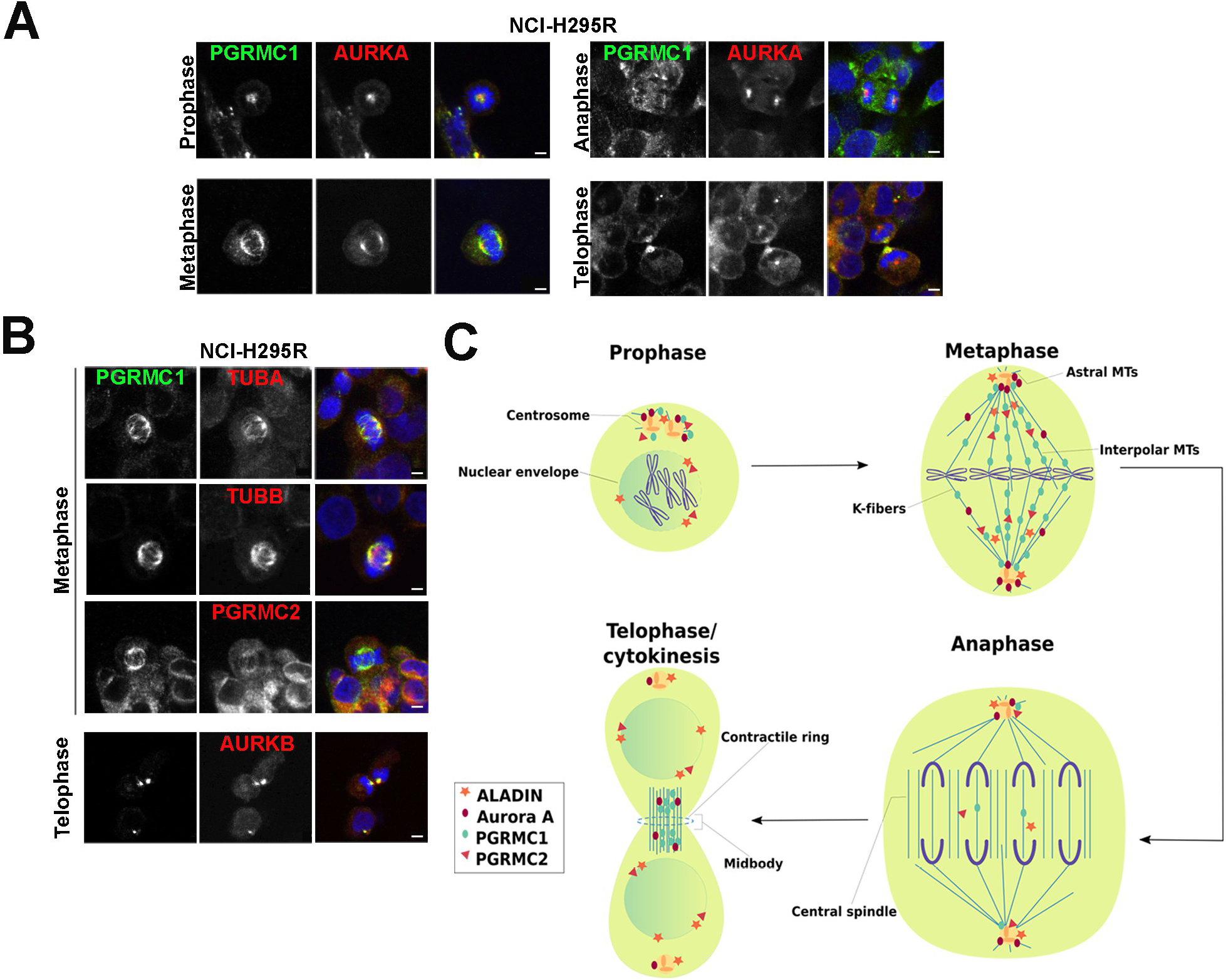
PGRMC1 is restricted to the mitotic spindle and to the mid-body during cytokinesis. (**A**) Human adrenocortical NCI-H295R cells at different cell division stages were stained with anti-PGRMC1 (green), anti-Aurora kinase A (AURKA) (red) and DAPI (blue). (**B**) Human adrenocortical NCI-H295R cells at meta- and telophase were stained with anti-PGRMC1 (green), anti-α-tubulin (TUBA) (red), anti-β-tubulin (TUB) (red), anti-PGRMC2 (red), anti-Aurora kinase B (AURKB) (red) and DAPI (blue). Scale bars: 5 μm. (**C**) Schematic drawing of cellular localization of ALADIN, Aurora kinase A, PGRMC1 and PGRMC2 during mitosis. MT, microtubule.

PGRMC1 efficiently localizes with α- (TUBA) and β-tubulin (TUBB) to metaphase bipolar spindles in adrenal NCI-H295R cells (Figure 2B). It has already been documented that PGRMC1 interacts with β- tubulin that is one of the two main components of MTs which are built from heterodimers of α- and β-tubulin (Scholey *et al*., 2003; Lodde and Peluso, 2011). Heterodimers are arranged in a head-to-tail fashion into protofilaments whereby in humans 13 of such protofilaments laterally arrange into tubular structures of about 25 nm diameter (Meunier and Vernos, 2012). Within MTs α- and β-tubulin heterodimers reveal a distinct intrinsic polarity in which the minus-end confers to the end where α-tubulin is exposed and the plus-end where β-tubulin is exposed (Meunier and Vernos, 2012). Microtubule polymerization is done at the β-tubulin plus-end and initial nucleation of MTs is facilitated by γ-tubulin including a complex of several proteins (Meunier and Vernos, 2012). Microtubules undergo dynamic cycles of polymerization and depolymerization which is called dynamic instability and is achieved by a variety of regulatory proteins (Meunier and Vernos, 2012). Co-localization of PGRMC1 with β- and γ-tubulin has been proposed before, but a direct interaction of PGRMC1 could only be shown for β-tubulin (Lodde and Peluso, 2011). Here, we show that β-tubulin is a positive target for a novel interaction with PGRMC1 and that association of PGRMC1 with β-tubulin possibly influences polymerization at plus-ends of MTs during mitosis. Furthermore, we show that PGRMC2 was loosely to the mitotic spindle in adrenal metaphase cells and weakly co-localized with PGRMC1 during mitosis (Figure 2B). In interphase cells we previously reported that PGRMC2 co-localizes with the nucleoporin ALADIN to the perinuclear space (Jühlen *et al*., 2016). Here, it appeared that PGRMC2 like ALADIN diffuses loosely in mitotic cells and that only PGRMC1 could be efficiently restricted to the mitotic bipolar spindle (schematically summarized in Figure 2C). It is known that PGRMC1 and PGRMC2 interact to suppress entry into cell cycle (Peluso *et al*., 2014). With immunofluorescence we could however not target PGRMC2 to the same site as PGRMC1 during mitosis. Thus, we hypothesize that PGRMC2 plays a distinct role in regulating the function of PGRMC1 at the mitotic spindle and presumably during cell division.

### Loss of ALADIN results in mis-localization of PGRMC1 and Aurora kinase A during metaphase

We have previously shown that the nucleoporin ALADIN interacts with microsomal PGRMC2 and that ALADIN is a new important co-factor of mitotic controller Aurora kinase A (Carvalhal *et al*., 2015; Jühlen *et al*., 2016). It is thought that PGRMC2 binds its protein homologue PGRMC1 resulting in suppression of cell cycle (Peluso *et al*., 2014). Here, we demonstrate that over-expression of ALADIN, PGRMC1 or PGRMC2 in adrenal cells leads to impaired cellular proliferation (Figure 1B), and that PGRMC1 is restricted to the mitotic bipolar spindle (Figure 2). Since both ALADIN and PGRMC2 loosely diffuse in mitotic cells compared to PGRMC1 (Figure 2C), we assume that ALADIN and PGRMC2 possibly exploit a regulatory role during cell division. Elucidating this novel role for ALADIN during cell division shall establish new avenues in explaining parts of the pathogenesis in triple A syndrome. We recently showed that a depletion of ALADIN in triple A patient fibroblasts affects the localization of PGRMC2 at the perinuclear ER (Jühlen *et al*., 2016). Thus, we now tested the effect of over-expression or depletion of ALADIN on the localization of PGRMC1 and Aurora kinase A. We used stable GFP-ALADIN over-expressing and inducible *AAAS* knock-down adrenocortical cells as well as skin fibroblasts from triple A patients. For this purpose, we chose patient cells with three different, and *in vivo* frequently occurring homozygous mutations in the *AAAS* gene: a donor splice mutation IVS14+1G>A (IVS14), a point mutation c.787T>C leading to the missense mutation S263P and a point mutation c.884G>A leading to the nonsense mutation W295X.

Using immunofluorescence we revealed that stable GFP-ALADIN over-expressing adrenocortical NCI-H295R cells failed to correctly localize Aurora kinase A to metaphase spindle poles (Figure 3, A and B, top panels). Aurora kinase A was not restricted to spindle poles in these cells but spread outward onto MT spindles compared to control cells over-expressing GFP (Figure 3, A and B, top panels). The same phenotype could be seen in inducible adrenocortical NCI-H295R1-TR cells depleted for ALADIN compared to control cells expressing scrambled shRNA (Figure 3, A and B, middle panels). Aurora kinase A normally resides at the centrosome and spindle poles where it regulates centrosome maturation and MT spindle assembly (Barr and Gergely, 2007). Carvalhal et al. reported that mitotic HeLa cells depleted for or overexpressing ALADIN have less Aurora kinase A at centrosomes and higher amounts of it at spindle fibers (Carvalhal *et al*., 2015). Our findings further prove that ALADIN partially regulates mitotic Aurora kinase A and raise the question why an equilibrated level of ALADIN protein is of great importance for proper localization of Aurora kinase A. Localization of PGRMC1 was altered in adrenocortical cells depleted for ALADIN (Figure 3, A and B, middle panels). PGRMC1 was still correctly targeted to the bipolar MT spindle but distribution onto spindle fibers did not fully extend onto chromosomal kinetochores compared to control scrambled shRNA-expressing cells (Figure 3, A and B, middle panels). Only depletion of ALADIN and not over-expression affected localization of PGRMC1 in mitotic cells. We know that ALADIN directly regulates Aurora kinase A and therefore, any change in level of ALADIN protein alters Aurora kinase A localization and function during mitosis (Carvalhal *et al*., 2015). As only depletion of ALADIN alters localization of PGRMC1 in immunofluorescence, it can be assumed that during mitosis ALADIN not directly regulates PGRMC1 but does so rather through secondary mechanisms.

**Figure 3.**
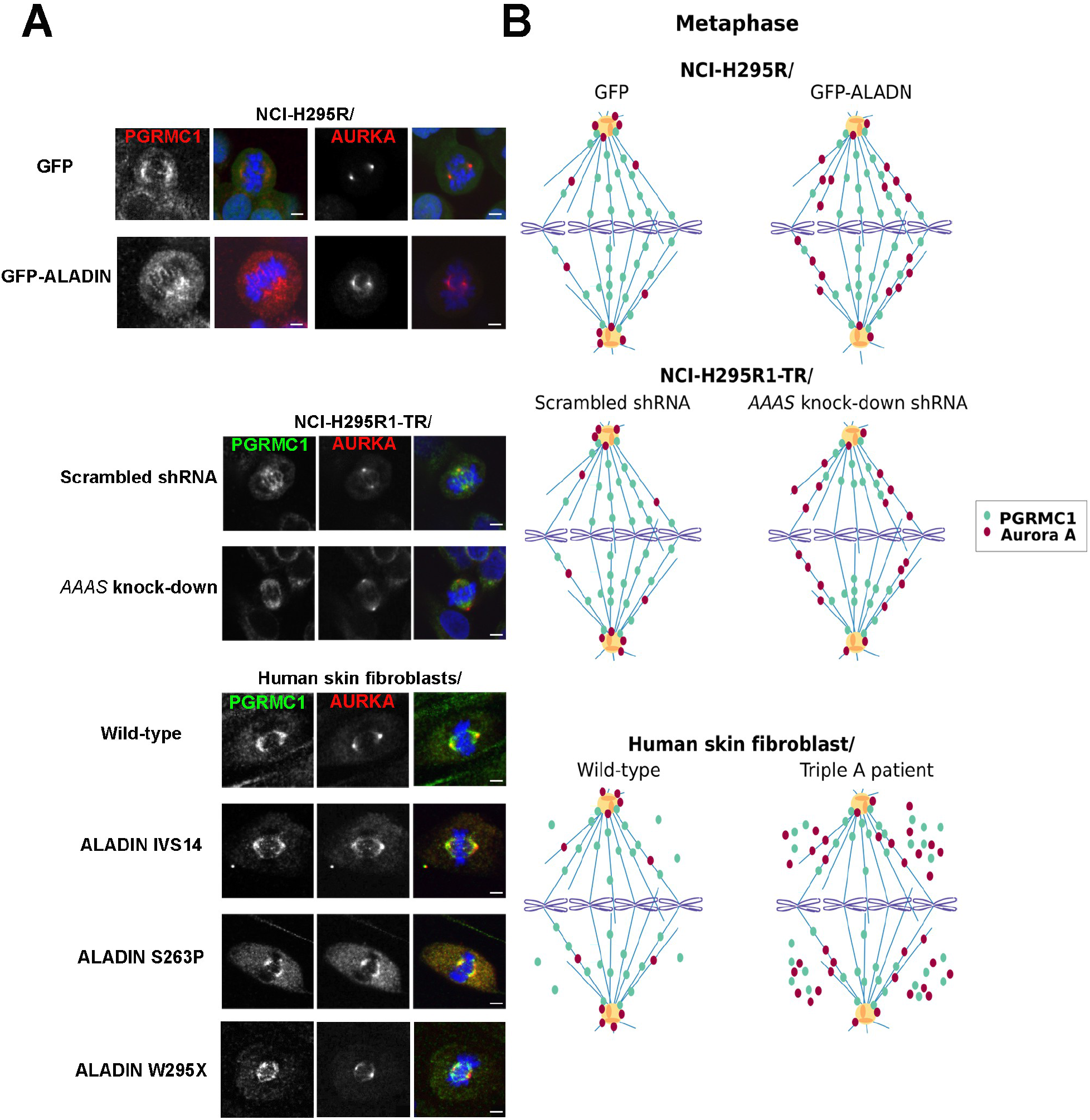
Loss of ALADIN mis-localizes PGRMC1 and Aurora kinase A during mitosis. (**A**) Human adrenocortical NCI-H295R GFP and GFP-ALADIN over-expressing cells, NCI-H295R1-TR scrambled shRNA and *AAAS* shRNA (*AAAS* knock-down) cells and human skin fibroblasts of healthy wild-type donors and triple A syndrome patients were stained with anti-PGRMC1 (green), anti-Aurora kinase A (AURKA) (red) and DAPI (blue). The different mutations in the human ALADIN protein are denoted as IVS14, S263P and W295X. Scale bars 5 μm, but for NCI-H295R GFP over-expressing cells: 10 μm. (**B**) Schematic drawing of the immunofluorescence staining of PGRMC1 and Aurora kinase A in (A).

Next, we tested our assumptions using fibroblasts from triple A patients. It has been described that triple A patient fibroblasts present with disorganized metaphase plates, shorter MT spindles and less active Aurora kinase A at spindle poles (Carvalhal *et al*., 2015). Here we show in patient fibroblasts that ALADIN depletion results in poleward spread of Aurora kinase A onto metaphase spindles and moreover, in accumulation in the cytoplasm (Figure 3, A and B, bottom panels). The highest accumulation of Aurora kinase A in the cytoplasm was seen in patient cells carrying the ALADIN missense mutation S263P and the nonsense mutation W295X (Figure 3, A and B, bottom panels). PGRMC1 was correctly restricted to the metaphase bipolar spindle with little decreased spread in direction of kinetochore chromosomes as seen in adrenocortical cells depleted for ALADIN (Figure 3, A and B, bottom panels). Nevertheless, PGRMC1 also accumulated in the cytosol of triple A fibroblasts with highest levels in patient cells carrying the missense mutation S263P and the nonsense mutation W295X (Figure 3, A and B, bottom panels). Thus, immunofluorescence results in triple A patient fibroblasts verified our previous findings and patient cells even presented with a more profound phenotype regarding Aurora kinase A and PGRMC1 mis-localization than adrenocortical ALADIN knock-down cells.

Our recent work presented that depletion of ALADIN alters the localization of PGRMC2 in triple A patient fibroblasts and leads to an increased level of PGRMC2 at the perinuclear ER (Jühlen *et al*., 2016). Furthermore, we showed that adrenal tissue of female ALADIN knock-out mice exploits higher levels of PGRMC2 protein compared to female wild-type animals (Jühlen *et al*., 2016). Here we show that fibroblasts from triple A patients carrying the ALADIN missense mutation S263P or the nonsense mutation W295X had about two-fold increased levels of PGRMC2 on mRNA and protein level compared to anonymized healthy control fibroblasts (Figure 4, A and B). Cells from the patient carrying the donor splice mutation IVS14 presented also with increased protein levels of PGRMC2 but quantitative real-time PCR data was not significant (Figure 4, A and B). We hypothesize that increased levels of PGRMC2 in fibroblasts from triple A patients result from the loss of ALADIN in these cells. Furthermore, we assume since ALADIN is a novel interactor of PGRMC2 that ALADIN negatively regulates microsomal PGRMC2. A direct or indirect negative regulation of PGRMC2 results in accumulation of PGRMC2 upon ALADIN depletion. We revealed that ALADIN depleted patient fibroblasts fail to correctly target PGRMC1 fully to the mitotic spindle fibers, instead PGRMC1 accumulates in the cytosol. Levels of PGRMC1 mRNA were not altered in these cells (**Figure S1**) and we already presented that over-expression of either PGRMC1 or PGRMC2 impairs cellular proliferation (Figure 1B). Based on the finding that PGRMC1 interacts with PGRMC2, we think that increased levels of PGRMC2 displace PGRMC1 from its correct localization at the mitotic spindle and target PGRMC1 with PGRMC2 to the cytosol impairing cell division and proliferation. Over-expression of PGRMC1 alone would probably also result in the same phenotype while targeting high amount of PGRMC1 to the cytoplasm.

**Figure 4.**
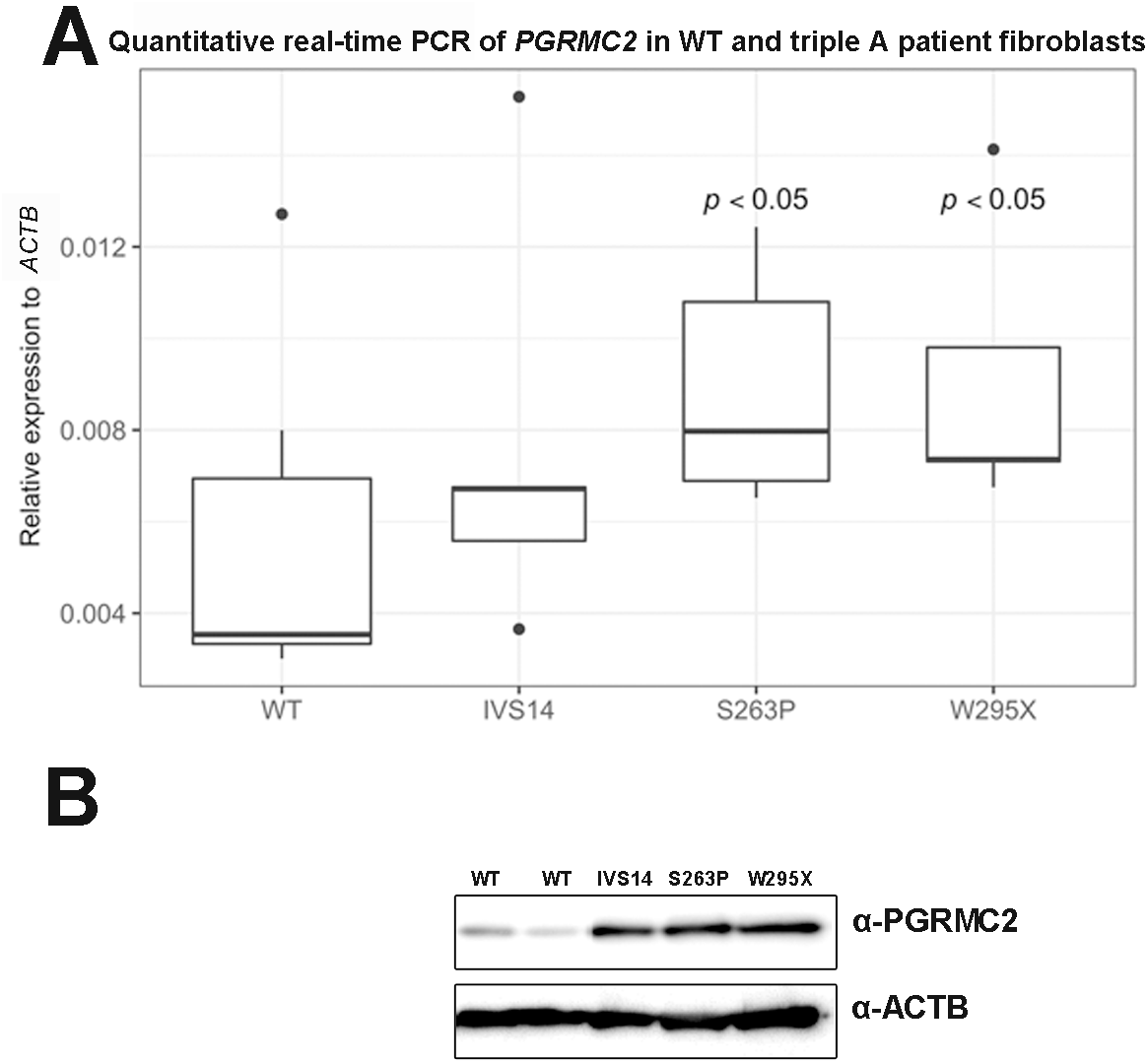
PGRMC2 is over-pressed in triple A patient fibroblasts. (**A**) Total RNA was isolated from human skin fibroblasts of healthy wild-type donors and patients with triple A syndrome. The different mutations in the human ALADIN protein are denoted on the x-axis of the diagram: IVS14, S263P and W295X. WT, wild-type. Significant differences were measured with unpaired Wilcoxon–Mann–Whitney U-test. Boxplot widths are proportional to the square root of the samples sizes. Whiskers indicate the range outside 1.5 times the inter-quartile range (IQR) above the upper quartile and below the lower quartile. Outliers were plotted as dots. (**B**) Total protein was isolated from human skin fibroblasts of healthy wild-type donors and triple A patients followed by western blot with indicated antibodies.

Adrenocortical ALADIN knock-down cells did not present with alteration of *PGRMC2* expression (Jühlen *et al*., 2016). Additionally, in immunofluorescence during mitosis PGRMC1 and Aurora kinase A were not targeted to the cytosol in these cells (Figure 3, A and B, middle panels). Moreover, fibroblasts from the patient carrying the donor splice mutation IVS14 had less PGRMC1 and Aurora kinase A targeted to the cytosol (Figure 3, A and B, bottom panels). Future research has to address in more detail whether the loss of regulation of PGRMC2 and Aurora kinase A is dependent on different levels in ALADIN protein and on different kinds of mutations in the *AAAS* gene. With our previous work we have given evidence, that depletion of ALADIN impairs mitotic cell division and can explain parts of the pathogenesis of triple A syndrome. Our new results shall be the basis for more extended research focusing on mis-regulated PGRMC2 and Aurora kinase A due to loss of ALADIN.

### PGRMC1 localizes to microtubule (MT) kinetochore fibers

The mature bipolar mitotic spindle is made from different subclasses of MTs depending on their position, functionality and organization: astral MTs, interpolar MTs and kinetochore-fibers (K-fibers) (Figure 1C) (Meunier and Vernos, 2012). Astral MTs connect the centrosome with the cell cortex and regulate centrosome separation and spindle positioning (Rosenblatt, 2005). However, it has been shown that mitosis can occur without astral MTs (Khodjakov *et al*., 2000; Mahoney *et al*., 2006). The main body of the mitotic spindle is made up by dynamic interpolar MTs. Interpolar MTs originate from the centrosome towards the center of the spindle and connect in an anti-parallel fashion with interpolar MTs originating from the opposite spindle pole (Meunier and Vernos, 2012). Some interpolar MTs however are shorter and do not emanate have way through the mitotic spindle (Mastronarde *et al*., 1993). Interpolar MTs are the main component of the central spindle during anaphase and maintain spindle polarity and chromosome segregation (Glotzer, 2009; Vanneste *et al*., 2011). However, interpolar MTs only indirectly establish chromosome segregation because this function is facilitated by K-fibers (Meunier and Vernos, 2012). K-fibers are big bundles of 20-40 parallel MTs and are responsible of chromosomes attachment to spindle poles and facilitate sister chromatid segregation (McEwen *et al*., 1997; Meunier and Vernos, 2012). K-fibers are less dynamic and therefore have an average half-life of 4-8 min comparable to that of interphase MTs with 9-10 min (Bakhoum *et al*., 2009; Meunier and Vernos, 2012). The average half-lifes of astral and interpolar MTs are less than 1 min (Zhai *et al*., 1995; Meunier and Vernos, 2012). Therefore, K-fibers are the most stable MTs when exposed to cold or depolymerizing agents like nocodazole.

From previous work we know that a loss of ALADIN negatively affects K-fiber stability (Carvalhal *et al*., 2015). Moreover, Lodde and Peluso showed that PGRMC1 affects microtubule stability and mitotic progression by the action of progesterone (Lodde and Peluso, 2011). As PGRMC1 localizes to the metaphase bipolar spindle and ALADIN depletion through PGRMC2 up-regulation negatively affects the targeting of PGRMC1 to the spindle, we were eager to find out whether PGRMC1 localizes to a distinct subclass of MTs during mitosis. We exposed cells before immunostaining to cold and observed whether PGRMC1 was still targeted to the spindle in mitotic cells. A positive result would exclude a localization of PGRMC1 to astral or interpolar MTs and would emphasize a role of PGRMC1 at K-fibers. Furthermore, we targeted known K-fiber proteins during immunofluorescence and observed the localization of PGRMC1 compared to these. These K-fiber proteins were firstly, TACC3 (also known as maskin) which is together with clathrin and chTOG (also known as CKAP5) responsible for K-fiber bundling and secondly NDC80 which belongs to the KMN complex (KNL1-MIS12-NDC80 complex) that attaches K-fiber plus-ends to the outer chromosomal kinetochore through polymerization and de-polymerization (Joglekar *et al*., 2010; Booth *et al*., 2011). In order to visualize the MT spindle we also stained for α-tubulin (TUBA) and to observe the centromeric region we stained for CENPB (Ando *et al*., 2002). We used adrenocortical NCI-H295R cells and additionally verified our results in a different cell type using human skin fibroblasts from healthy donors and from earlier mentioned triple A patients.

In Figure 5, A and B, we well document that after cold treatment PGRMC1 was still restricted to MT spindles in adrenocortical NCI-H295R cells and human skin fibroblasts. α-tubulin still co-localized with PGRMC1 and thus, targeted PGRMC1 to cold-stable K-fibers (Figure 5, A and B). Furthermore, immunofluorescence of CENPB marked the centromeric region of the condensed chromosomes containing the kinetochore. It can be seen that PGRMC1-positive spindle K-fibers reached out to the centromeric region being attached by the KMN complex visualized by immunostaining of NDC80 (Figure 5, A and B). Immunofluorescence of PGRMC1 further partly co-localizes with staining of the MT-bundling protein TACC3 in adrenocortical cells and skin fibroblasts (Figure 5, A and B). In Figure 5C the same immunofluorescence staining pattern can be seen in cold-treated skin fibroblasts from triple A patients carrying the donor splice mutation IVS14, the missense mutation S263P or the nonsense mutation W295X. However, these cells present with shorter and more diffuse K-fibers compared to fibroblasts from healthy donors (Figure 5, B and C). We hypothesize that this observation is due to the previously documented decreased stability of K-fibers upon ALADIN depletion (Carvalhal *et al*., 2015).

**Figure 5.**
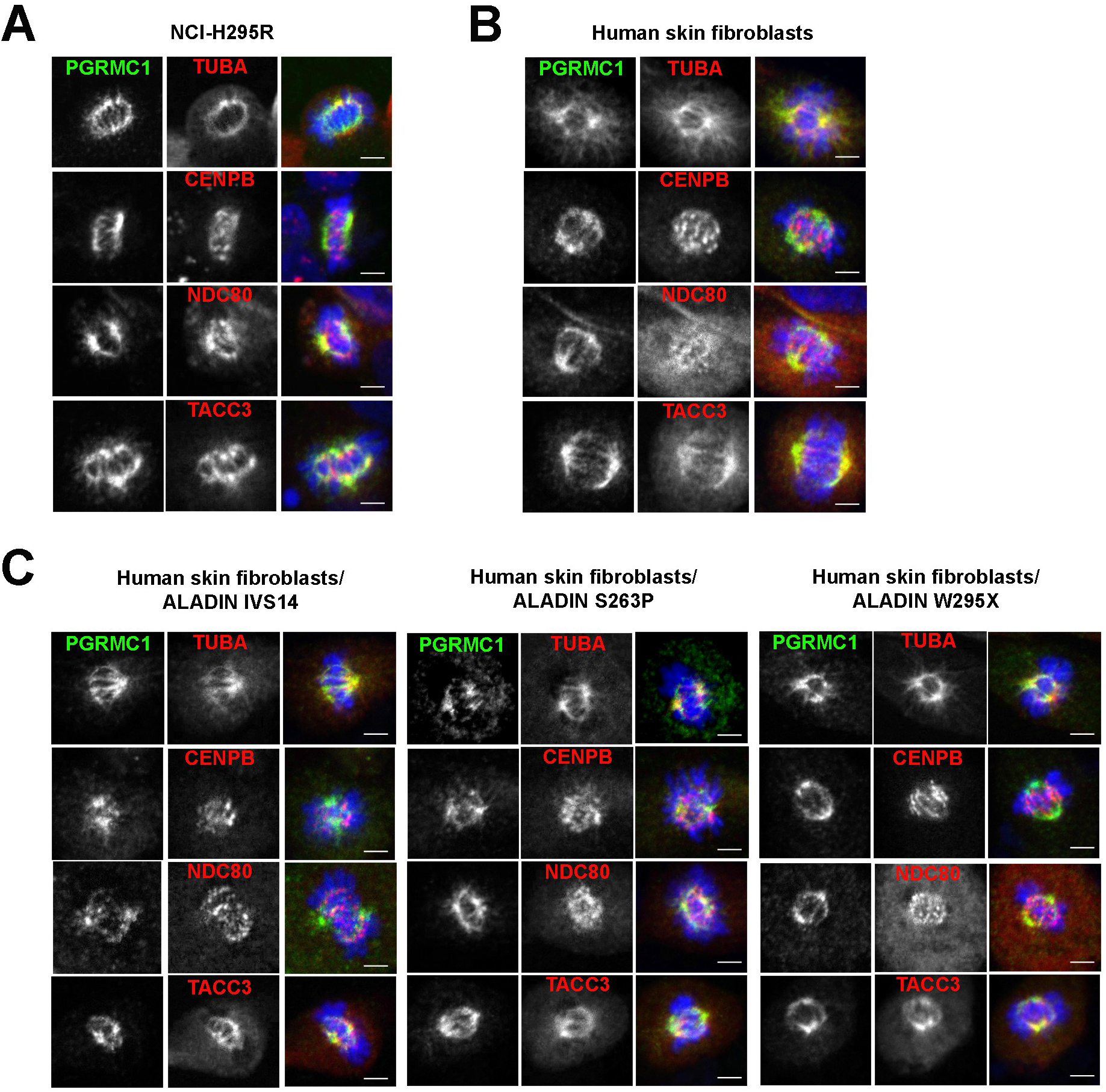
PGRMC1 localizes to cold-stable kinetochore fibers. Cells were cold-treated and stained with anti-PGRMC1 (green), anti-α-tubulin (TUBA) (red), anti-CENPB (red), anti-NDC80 (red), anti-TACC3 (red) and DAPI (blue). (**A**) Human adrenocortical NCI-H295R cells. (**B**) Human skin fibroblasts of healthy wild-type donors. (**C**) Human skin fibroblasts of triple A patients. The different mutations in the human ALADIN protein are denoted as IVS14, S263P and W295X. Scale bars 5 μm.

In conclusion we document here that PGRMC1 localizes to cold-stable MT K-fibers in human adrenocortical cells and skin fibroblasts. We verify that K-fibers are less stable upon ALADIN loss of function in triple A patient fibroblasts and we present that these cells fail to target PGRMC1 efficiently to the mitotic spindle. Thus, we assume that part of the phenotype seen in triple A patient cells can be caused by a mis-regulated interaction between ALADIN and PGRMC2 resulting in mis-localization of PGRMC1 during mitosis and less stable K-fibers. Further studies are required to address the exact function of the microsomal protein PGRMC1 at K-fibers and to solve the question whether it is involved in MT bundling and/or MT maturation.

## MATERIALS AND METHODS

### Cell culture

All adrenal carcinoma NCI-H295R cells were cultured in DMEM/F12 medium (Lonza, Cologne, Germany) supplemented with 1 mM L-glutamine (Lonza, Cologne, Germany), 5% Nu-serum (BD Biosciences, Heidelberg, Germany), 1% insulin-tranferrin-selenium) (Gibco, Life Technologies, Darmstadt, Germany) and 1% antibiotic-antimycotic solution (PAA, GE Healthcare GmbH, Little Chalfont, United Kingdom).

NCI-H295R cells stably expressing GFP-ALADIN fusion protein or GFP were generated as described previously using the gamma-retroviral transfer vectors pcz-CFG5.1-GFP-AAAS and pcz-CFG5.1-GFP (Kind *et al*., 2009).

NCI-H295R1-TR cells stably expressing *AAAS* shRNA (*AAAS* knock-down) or scrambled shRNA were generated by our group as previously reported (Jühlen *et al*., 2015). These cells were cultured with 100 μg/ml zeocin (InvivoGen, Toulouse, France) supplemented in culture medium. Doxycyline hydrochloride (MP Biomedicals, Eschwege, Germany) was used at 1 μg/ml for 48 h to turn on the expression of the shRNA sequence.

Triple A patient skin fibroblasts and human anonymized control skin fibroblasts were obtained and cultured as described earlier (Kind *et al*., 2010). All fibroblasts were cultured until passage 20 at the most. Informed consent was obtained from all subjects and experiments were approved by the local ethics review board (Medical Faculty, Technische Universität Dresden, EK820897).

### Transient adrenal cell transfection

Cells were cultured for proliferation analysis in 24-well culture dishes at a density of 0.4 x10^5^ cells/well or for immunofluorescence microscopy onto cover slips (Carl Zeiss, Jena, Germany) in 6-well culture dishes at a density of 1.6 × 10^5^ cells/well 24 h before subsequent transfections. Cells were transfected using X-treme GENE HP DNA transfection reagent (Roche Diagnostics, Mannheim, Germany). The plasmids for transient transfections pEGFP-C1-PGRMC1 and pCMV6-AC-PGRMC2-GFP vector (RG204682) (OriGene Technologies, Rockville MD, USA) were used at a concentration of 0.01 μg/μl at an optimized transfection ratio of 1:4 diluted in pure DMEM/F12. Proliferation was monitored after 24 h after transfection. Cells for immunofluorescence were fixed after 48 h.

### Proliferation analysis

Cells were seeded in 24-well culture dishes at a density of 0.4 x10^5^ cells/well 24 h before proliferation analysis. Confluence measurement was done using live cell imaging on IncuCyte Zoom (Essen BioScience, Ann Arbor MI, USA) over at least 65 h. Measurement was done at least in triplicate and experiments were repeated at least twice.

### Immunofluorescence microscopy

Cells grown on glass cover slips were fixed with 4% PFA in PBS for 5 min, permeabilized with 0.5% Triton-X-100 in PBS for 5 min and then fixed again. In order to assess K-fiber microtubule stability, cells were put on ice for 10 min prior fixation and permeabilization as reported elsewhere (Carvalhal *et al*., 2015). Blocking was performed with 2% BSA/0.1% Triton-X-100 in PBS for 30 min at room-temperature (RT).

All antibodies used for immunofluorescence were diluted in blocking solution. Primary antibodies anti-ALADIN (B-11: sc-374073) (1:25), anti-AURKA (C-1: sc-398814) (1:50), anti-AURKB (A-3: sc-393357) (1:50), anti-CENPB (F-4: sc-376283) (1:50), anti-NDC80 (C-11: sc-515550) (1:50), anti-PGRMC1 (C-4: sc-393015) (1:50), anti-PGRMC2 (F-3: sc-374624) (1:50), anti-TACC3 (C-2: sc-376883) (1:50), anti-TUBA (DM1A: sc-32293) (1:50) and anti-TUBB (D-10: sc-5274) (1:50) (Santa Cruz Biotechnology Inc., Heidelberg, Germany) were incubated at 4°C over-night in a humidified chamber. Secondary antibodies Alexa Fluor 488 goat anti-mouse IgM and Alexa Fluor 568 goat anti-mouse IgG (1:500) (Molecular Probes, Life Technologies) were incubated one hour at RT in the dark. Excess antibodies after primary and secondary antibody staining were removed by three washing steps using 0.1% Triton-X-100 in PBS for 5 min. Cover slips were mounted onto microscope slides with VECTASHIELD mounting medium for fluorescence with DAPI (Vector Laboratories, Burlingame, CA, USA).

Fluorescence was imaged using the confocal laser scanning microscope Zeiss LSM 510 with Zeiss EC Plan-Neofluar 40x objective/ 1.3 Oil and the following lasers: diode 405 nm, Argon 488 nm and DPSS 561 nm (Carl Zeiss). Images were acquired and processed using equipment of the Core Facility Cellular Imaging at the Medical Theoretical Centre in Dresden. The experiments were repeated at least three times.

### RNA extraction, cDNA synthesis and quantitative real-time PCR

Total RNA from cultured cells was isolated using the NucleoSpin RNA (Macherey-Nagel, Düren, Germany) according to the protocol from the manufacturer. Purity of the RNA was assessed using Nanodrop Spectrophotometer (ND-1000) (NanoDrop Technologies, Wilmington DE, USA). The amount of 500 ng of total RNA was reverse transcribed using the GoScript Reverse Transcription System (Promega, Mannheim, Germany) following the protocols from the manufacturer. Primers for the amplification of the target sequence were designed using Primer Express 3.0 (Applied Biosystems, Life Technologies) and compared to the human genome database for unique binding using BLAST search (National Center for Biotechnology Information, U.S. National Library of Medicine, 2013). Primers for *PGRMC1* (forward, reverse and probe) were used as previously described (Hlavaty *et al*., 2016). The primer sequences are listed in the supplementary data of this article (**Table S1**).

The qPCR amplifications were performed in triplicates using the GoTaq Probe qPCR Master Mix (Promega) according to the manufacturer’s reaction parameter on an ABI 7300 Fast Real-Time PCR System (Applied Biosystems, Life Technologies). In all results repeatability was assessed by standard deviation of triplicate Cts and reproducibility was verified by normalizing all real-time RT-PCR experiments by the Ct of each positive control per run. The experiments were repeated at least five times.

### Immunoblots

After SDS-PAGE separation onto 4-12% PAGE (150 V for 1.5 h) and electroblotting (30 V for 1.5 h) (Invitrogen, Life Technologies) onto Amersham hybond-ECL nitrocellulose membrane (0.45 μm) (GE Healthcare GmbH, Little Chalfont, United Kingdom) non-specific binding of proteins to the membrane was blocked by incubation in PBS containing 3% BSA at RT.

The membrane was then probed with primary antibodies either anti-ACTB (clone AC-74) (1:40000 in 3% PBS/BSA) (Sigma-Aldrich, Munich, Germany), anti-PGRMC1 (C-4: sc-393015) (1:100 in 3% PBS/BSA) or anti-PGRMC2 (F-3: sc-374624) (Santa Cruz Biotechnology, Inc.) (1:100 in 3% PBS/BSA) over-night at 4°C. Secondary antibodies goat anti-mouse IgG conjugated to horseradish peroxidase (1:5000 in 3% PBS/BSA) (Cell Signalling Technology Europe B.V., Leiden, Netherlands) were incubated one hour at RT. Protein bands were detected using ECL system and visualized on autoradiography film (Hyperfilm ECL; GE Healthcare, Munich, Germany).

### Statistics

Statistical analyses were made using the open-source software R version 3.4.2 and R Studio version 1.0.136 (R Core Team, 2017). Unpaired Wilcoxon-Mann-Whitney U-test was performed. During evaluation of the results a confidence interval alpha of 95% and P values lower than 0.05 were considered as statistically significant. Results are shown as box plots which give a fast and efficient overview about median, first and third quartile (25^th^ and 75^th^ percentile, respectively), interquartile range (IQR), minimal and maximal values and outliers.

Growth curve analysis and growth constant k (slope of linear regression line) calculation was done using multilevel regression technique using R Studio.

## ACKNOWLEDGEMENTS

We thank Waldemar Kanczkowski (Technische Universität Dresden, Dresden, Germany) for providing the NCI-H295R cells. Barbara Kind (Technische Universität Dresden, Dresden, Germany) generously generated pseudo retroviruses containing pcz-CFG5.1-GFP-ALADIN and pcz-CFG5.1-GFP. Alexandra Wendler (Universität Heidelberg, Mannheim, Germany) kindly provided pEGFP-C1-PGRMC1 plasmid.

## Abbreviations

ALADIN: alacrima-achalasia-adrenal insufficiency neurologic disorder
CYP: cytochrome P450
ER: endoplasmic reticulum
GFP: green fluorescent protein
NADPH: Nicotinamide adenine dinucleotide phosphate
NUP: nucleoporin
PGRMC1 and 2: progesterone receptor membrane component 1 and 2

